# Evaluating First-Pass, High Protein Capacity Desalting Techniques For Phosphoproteomics Applications

**DOI:** 10.1101/2025.06.03.657744

**Authors:** Aurora Callahan, Aisharja Mojumdar, Arthur R. Salomon, Nicholas A. DaSilva

## Abstract

Many commercial desalting products exist for pre-MS peptide cleanup, although few exist that can handle the high protein input (≥ 4 mg) required for phosphotyrosine enrichment. For these desalting products, the technical aptitude required for effective and organized desalting is often a barrier to entry for new users. Here, we evaluate four commercially available desalting techniques with varying degrees of automation, operational organization, and chemistries to determine the most cost-effective, user-friendly, and sensitive technique for protein profiling and phosphotyrosine (pY) enrichment. We find that TECAN Narrow Bore Extraction (NBE) products are the most cost effective per sample and least difficult to use, whereas ProtiFi’s S-Trap are the most expensive per sample and Pierce C18 spin columns have the worst operational organization. ProtiFi S-Trap vastly outperforms other desalting methods for peptide sequencing and protein profiling applications, uniquely identifying 25,654 unique peptide sequences and 375 unique proteins. Consistently, ProtiFi S-Trap samples show the deepest pY sequencing after Src SH2 superbinder enrichment, leading to the highest identification of significantly changing, biologically relevant pY sites in a Jurkat T cell signalling model. Our data show that ProtiFi S-Trap columns provide high peptide recovery, thus increasing meaningful pY site identification.

**Graphical Abstract:** 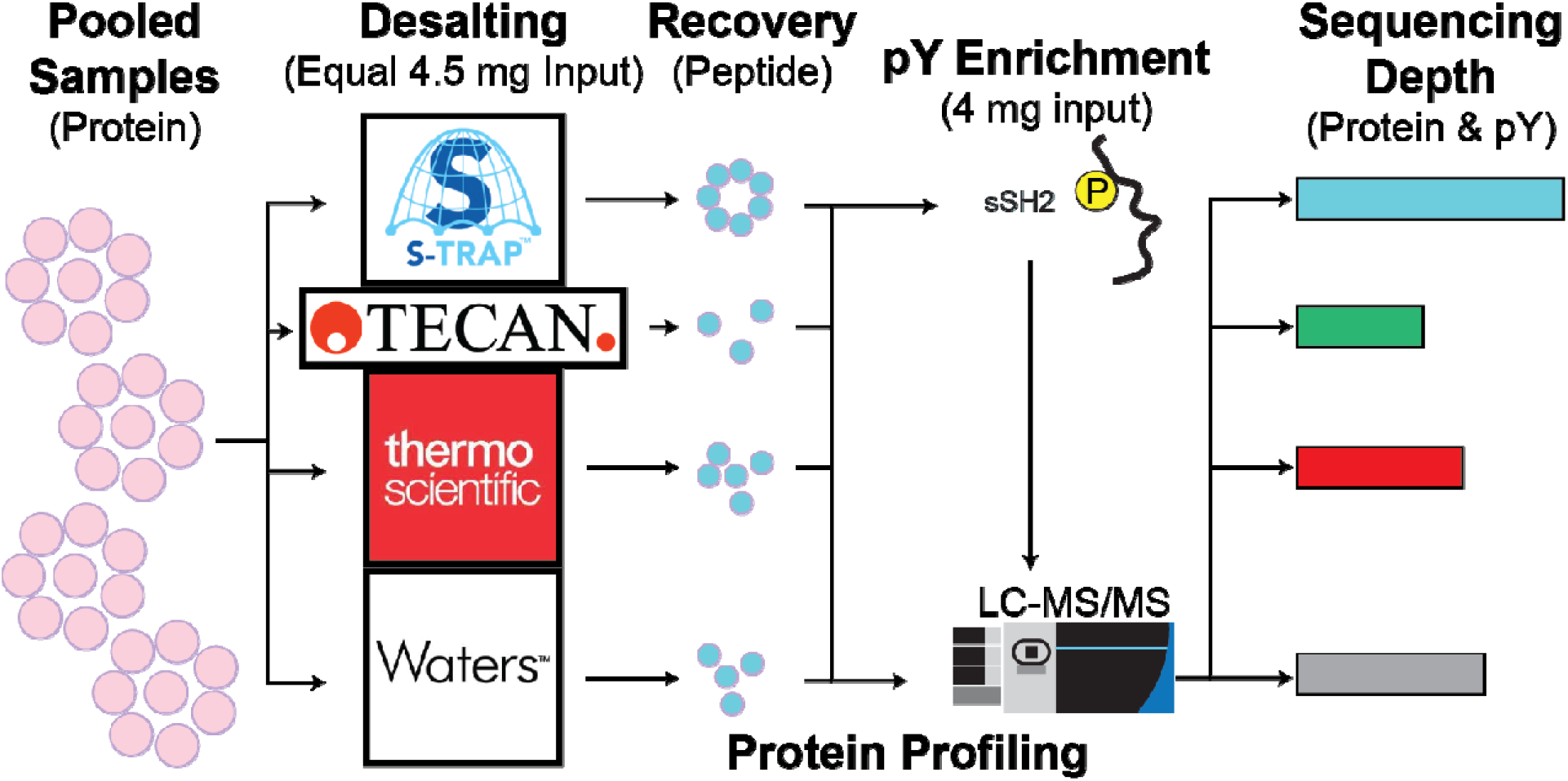

## 1 Introduction

Mass spectrometry-based proteomics is a rapidly evolving field in the biological sciences, with a skyrocketing number of users attempting proteomics projects, with users submitting more than 4,000 projects to the <u>PRIDE repository</u> per year between 2020-2023 and 3,241 in 2024^1^.

Improvements to MS instrumentation, methodology (bottom-up, top-down, and middle-down proteomics), and data processing techniques have increased the number of MS-based applications, including ultra deep shotgun protein profiling^2,3^, single-cell characterisation^4^, post-translational modification (PTM) enrichment^5,6^, interactomics^7^, structural proteomics^8^, spatial proteomics^9^, and top-down proteomics^10^. These applications have led to a wave of discoveries concerning animal and human disease state proteomes^11^ while generating new hypotheses and research aims for mechanistic biologists and translational researchers alike^12^. MS-based proteomics, sample preparation, nano-LC systems, and MS instrumentation are increasingly complex, requiring expertise to understand and implement, which acts as a barrier to entry to the field. Some users elect to use MS core facilities^13,14^ or collaborate with expert laboratories, while using commercially available sample preparation and enrichment products to generate contaminant-free peptides as part of common bottom-up proteomic workflows.

For bottom-up shotgun proteomics, the most common sample preparation method involves lysis/sonication (Urea or non-ionic detergents), denaturation, reduction (e.g dithiothreitol) alkylation (e.g iodoacetamide), and enzymatic protein digestion (Trypsin or Lys-C alone or in combination), desalting using reverse phase column chromatography (typically C18), and concentrating peptides via lyophilization or chilled centrivap^15^. Protocols for this style of sample preparation are well established, and are in the process of being automated at several levels using liquid handling (LiHa) systems. These methods, while requiring some knowledge of liquid handling machines, reduce the propensity for user error in classic sample preparation techniques, such as vacuum manifold or centrifugal desalting^16–18^. Despite this, many research groups and core facilities preferentially use classic sample preparation methods, particularly for the desalting step. Specific applications that require large protein input (∼5 mg per sample), such as PTM enrichment proteomics, suffer from a general lack of commercial desalting products, with even fewer products being automation capable.

Here, we evaluate four first pass, high input desalting products for use in protein profiling and phosphotyrosine enrichment applications using a Jurkat T cell signalling model. The products chosen (Waters Sep-Pak C18 vacuum cartridges, Pierce C18 spin columns, TECAN WWP2 Narrow Bore Extraction reverse phase columns, and ProtiFi S-Trap spin columns) are chosen based on their availability, chemistry, and methodology to represent a wide variety of commonly used desalting techniques. We find that Sep-Pak vacuum desalting is the most procedurally difficult, while Pierce C18 spin columns are the most organizationally challenging product to use. ProtiFi S-Trap spin columns provide the highest overall peptide recovery (∼75%), whereas TECAN WWP2 NBE columns provide the lowest variability yet lowest peptide recovery (∼30%). The higher percent recovery for ProtiFi S-Trap preparations correspond to significantly more peptide and protein identifications, with 25,653 peptides and 375 proteins identified exclusively in at least one ProtiFi S-Trap sample. ProtiFi S-Trap samples also showed the deepest pY site sequencing after Src SH2 superbinder pY enrichment, identifying ∼400 unique pY sites per sample, more than 700 unique pY sites total, and more than 400 unique pY sites exclusively in at least one ProtiFi S-Trap sample. Finally, deep pY sequencing in ProtiFi S-Trap samples increases the identification of significantly T cell receptor responsive pY sites on biologically meaningful proteins.

## 2 Materials and Methods

### 2.1 Cell culture and stimulation

Jurkat (clone E6.1) were maintained in RPMI 1640 with 10% FBS (Cytiva #SH30910.03), 1x penicillin-streptomycin-glutamine (HyClone #SV30082.01), 2.5 μg/mL Plasmocin (InvivoGen #ant-mpp) in a humidified incubator at 37 °C, 5% CO_2_. For stimulation, 4e8 cells per replicate (4e9 cells total) were collected, washed once in 1x DPBS, then resuspended at a concentration of 2e8 cells/mL in 1x DPBS and rested for 30 minutes at 37°C. A schematic of the sample preparation process is shown in Figure 1A. T cell receptor (TCR) stimulation was induced by the addition of α-CD3e (clone OKT3; Thermo #16-0037-85) and α-CD28 (clone CD28.6; Thermo #16-0288-81) for 30 seconds before addition of α-Mouse IgG (Jackson ImmunoResearch #115-005-062). The final concentration of primary and secondary antibodies were 2 μg/mL and 22 μg/mL, respectively. Stimulation was stopped by addition of 9 M Urea Lysis Buffer (9 M Urea, 1 mM sodium orthovanadate, 1 mM sodium pyrophosphate, 1 mM β-glycerophosphate in 20 mM HEPES). The ‘0 minute’ samples were generated by addition of 9 M Urea Lysis Buffer, then subsequent addition of primary and secondary antibody solutions to assure no cell stimulation occurred. Lysates were sonicated at 70% amplitude for 30 seconds, once on ice and centrifuged at 1,800 xg for 15 minutes before protein quantification by BCA (Thermo #23225).

**Figure 1.**
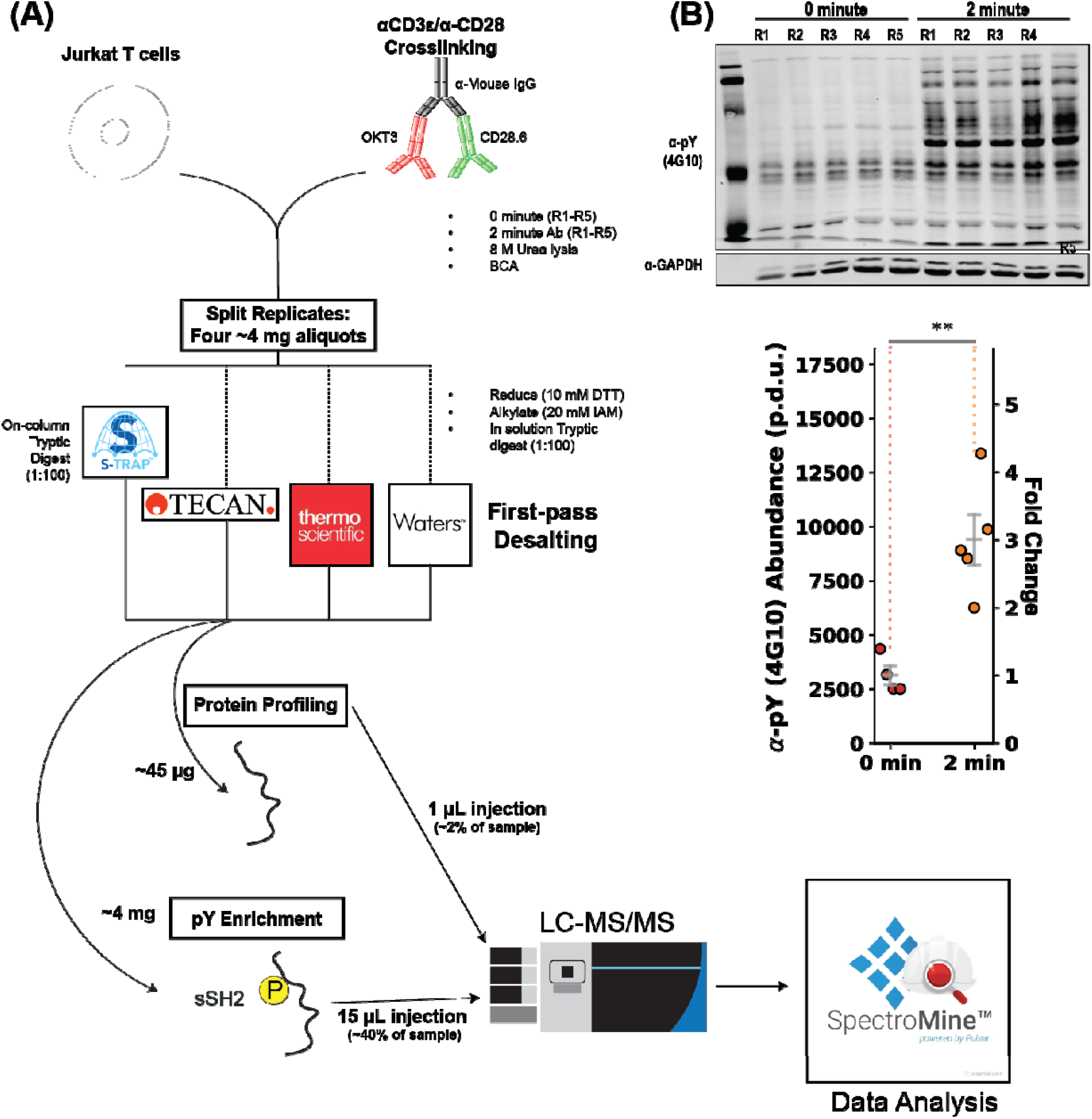
Workflow for evaluating high-protein input desalting products for proteomics applications. (A) Schematic representation of sample preparation. (B) α-pY (clone 4G10) Western blot showing pY induction after 2 minutes of α-CD3ε/α-CD28 crosslinking in Jurkat T cells with quantification. Error bars show S.E.M. ** indicates p<0.01 by a Welch’s T-test.

Western blotting samples were taken by diluting 50 μL from each sample 1:1 in 2x Lamelli Sample Buffer (4% SDS, 125 mM TRIS-HCl pH 6.8, 20% glycerol, 5% β-mercaptoethanol, 0.01% bromophenol blue). For each replicate, a solution containing approximately 4.5 mg of protein was allotted into four tubes, then volume-normalized to 2 mL using 4 M Urea and frozen before subsequent proteomics sample preparation.

### 2.2 Proteomics Sample processing

#### 2.2.1 Sep-Pak C18 vacuum desalting

Urea lysed samples were thawed and diluted in 20 mM HEPES to a volume of 4.45 mL before reduction using dithiothreitol (50 μL of 1 M DTT, final concentration 10 mM) at 37 C for 30 minutes and alkylation using iodoacetamide (500 μL 200 mM IAM, final concentration 20 mM) at room temperature in the dark for 30 minutes. Protein was digested overnight at 37 °C in solution using a 1:100 ratio of trypsin (Promega #V5113) to protein, as previously described^19–21^.

In the morning, samples were acidified to 1% trifluoroacetic acid (TFA) and centrifuged to remove debris before C18 vacuum desalting using Sep-Pak C18 cartridges, per the manufacturer’s instructions. Briefly, the vacuum apparatus was assembled as in Figure 1B and kept on except to discard waste and switch between waste tubes and collection tubes. Columns were equilibrated with 5 mL 100% acetonitrile (ACN) followed by two 3.5 mL washes in 0.1 % TFA. Samples were loaded (∼5.5 mL each), then washed in 1 mL, 5 mL, and 6 mL 0.1% TFA before elution in 7 mL of 40% ACN, 0.1% TFA (Final sample volume ∼7.5 mL). After elution, 1/100 (70 μL, diluted in 0.1% TFA) of each sample was taken for protein profiling. Samples were dried by overnight speedvac^19,20^.

#### 2.2.2 Pierce C18 spin column desalting

Overnight, in solution digestion was performed as in 2.2.1, acidified to 1% TFA, and centrifuged to remove debris the following morning. Samples were desalted using the Pierce C18 spin-columns (Thermo #89851) per the manufacturer’s instructions. Briefly, the caps of each column were labelled with the sample ID (Figure 1C), then spun at 5,000 xg for 1 minute to remove storage solution. Columns were equilibrated by centrifuging 300 μL of 100% ACN twice followed by 300 μL of 0.1% TFA twice at 5,000 xg for 1 minute each. About 5.5 mL of sample was loaded on each column by centrifuging 300 μL at a time at 3,000 xg for 1 minute each, discarding the flow through after each load (18 total loads). After sample loading, peptides were washed with 300 μL of 0.1% TFA twice by centrifugation at 3,000 xg for 1 minute each. After washing, peptides were eluted by centrifuging 300 μL of 50% ACN, 0.1% TFA twice at 3,000 xg for 1 minute. After elution, 1/100 (6 μL, diluted in 0.1% TFA) of each sample was taken for protein profiling. Samples were dried by overnight speedvac.

#### 2.2.3 TECAN A200 automated narrow bore extraction WWP2 reverse-phase desalting

Overnight, in solution digestion was performed as described previously for Pierce C18 and Waters products. Samples were acidified to 1% TFA, and centrifuged to remove debris the following morning. Samples were desalted using the Resolvex A200 (TECAN) positive pressure liquid handling system with TECAN Column NBE WWP2 1 mL 5.0 mg columns (TECAN #417-0051R-NBE). Narrow Bore Extraction columns were equilibrated with 900 μL of 80% ACN, 0.1% TFA then washed with 900 μL of 0.1% TFA using a 60 second positive pressure flash. Acidified samples were then loaded by sequential addition followed by a 45 second positive pressure flash, until the whole sample volume had passed through the column (6 loads). After loading, 900 μL of 0.1% TFA was passed through the columns with a 60 second flash followed by a 30 second flash. Samples were eluted into a collection plate with three sequential 45 second flashes of 200 μL 50% ACN, 0.1% TFA for a total elution volume of 600 μL. After elution, 1/100 (6 μL, diluted in 0.1% TFA) of each sample was aliquoted for protein profiling. Samples were dried by overnight speedvac.

#### 2.2.4 ProtiFi S-Trap sample preparation

In solution reduction and alkylation were performed by addition of 20 μL 1 M DTT (final concentration 10 mM) at 37 °C for 30 minutes, followed by 200 μL 200 mM IAM (final concentration 20 mM) at room temperature in the dark for 30 minutes. Samples were centrifuged to remove any debris before acidification to 0.1% TFA and 12 mL of sTrap buffer (90% Methanol, 100 mM TEAB, pH 7.1). Protein samples were bound to the ProtiFi sTrap midi columns (C02-midi-10) by sequential centrifuging of 4 mL sample solution through the columns at 3,000 xg for 5 minutes (no equilibration step necessary). Columns were washed with 4 mL of sTrap buffer three times before moving the column to a new centrifuge tube and adding digestion buffer (50 mM TEAB and 1:100 ratio of trypsin). The cap of the centrifuge tube was loosely shut, and tubes were wrapped in wet KimTech wipes before overnight trypsinisation at 37°C. The next morning, 500 μL of 50 mM TEAB was applied to the column and the digestion solution was centrifuged out of the column at 3,000 xg for 5 minutes. Next, 500 μL of 0.2% Formic acid was centrifuged through the column, followed by 50% ACN, 0.2% Formic acid to complete elution. After elution, 1/100 (20 μL, diluted in 0.1% TFA) of each sample was taken for protein profiling. Large volume samples were dried by 36-hour lyophilisation and small volume samples were dried by overnight speedvac.

### 2.3 hosphotyrosine enrichment with Src SH2 superbinder

Enrichment of tyrosine phosphorylated peptides was performed as previously described^19,20,22–24^. Briefly, ∼ 4 mg of peptides were resuspended in 1.4 mL of ice cold IAP buffer (10 mM sodium phosphate monobasic monohydrate, 50 mM sodium chloride, 50 mM morpholinopropanesulfonic) for ∼1 hour at room temperature with periodic vortexing. One hundred micrograms of Src SH2 superbinder (sSH2) conjugated agarose beads were allotted into 1.7 mL protein low-binding microcentrifuge tubes and washed 3 times in 1 mL of ice cold IAP buffer. sSH2 beads were incubated with samples for 2 hours at 4 °C on a 4 rpm rotator, and then centrifuged at 1,500 xg for 2 minutes. The supernatant was discarded, and the beads were washed 2 times in 1 mL of ice cold IAP buffer, then three times in 1 mL of ice cold HPLC grade water by centrifugation at 1,500 xg for 2 minutes at 4 C. After the last wash, all of the liquid was removed using an insulin syringe, and the peptides were eluted from the beads using 100 μL of 0.15% TFA on a 1,150 rpm shaker, twice. Peptide-containing solution was removed using an insulin syringe and desalted using Empore StageTips as described in Section 2.4.

### 2.4 StageTip desalting of enriched phosphopeptides

Enriched phosphopeptides were desalted using StageTips (Empore #70-2019-1001-3EA) per the manufacturer’s instructions. Tips were arranged in microcentrifuge tube adapters and activated by centrifuging 100 μL of 100% LC-MS grade methanol and 100 μL of 50% ACN, 0.1% TFA at 1,800 xg for 2 minutes. Tips were then washed by centrifuging 100 μL of 0.1% TFA twice at 1,800 xg for 2 minutes before loading samples and centrifuging at 1,800 xg for 2 minutes. Peptide-loaded tips were washed by centrifuging 100 μL of 0.1% TFA twice at 1,800xg for 2 minutes before elution by centrifuging 100 μL of 50% ACN, 0.1% TFA at 1,800 xg for 2 minutes, then reapplying the eluent solution to the column and centrifuging at 1,800 xg for 2 minutes. Eluent was then frozen and dried by speedvac before reconstitution in 5% Acetonitrile and 0.1% Formic acid and analysis by LC-MS.

### 2.5 Liquid chromatography tandem mass spectrometry

Whole cell digest samples were reconstituted in 50μL with 1μL injected into the LC/MS system. Enriched pY samples were reconstituted in 35μL with 15μL injected. Peptides were separated using a Vanquish Neo UHPLC on a 7cm long (2cm bed length) 3Lm particle size, 100 A, Acclaim PepMap C18 Trap column and IonOpticks Aurora Ultimate 25cm long x 75μm ID, 1.7μm particle size analytical column throughout a 60-minute gradient held at 300 nl/min. Solvent A consisted of Water with 0.1% Formic Acid and Solvent B consisted of 80% Acetonitrile and 0.1% Formic Acid.

The gradient consisted of loading/starting conditions of 5% B then to 30% B over 50 minutes followed by 95% B over 5 minutes then finishing at 99% B for the remaining 5 minutes. Combined control was used for loading and washing. Trap column was washed using 4 zebra washes, and the analytical column was washed using 3 alternating aqueous and organic cycles before being re-equilibrated for the next sample injection. Spray voltage was set to 2050V, Ion Transfer Tube Temperature was 275 °C, and a user-defined lock mass of 445.12003 was used. MS1 scans used wide quad isolation, orbitrap as the detector with a resolution of 120K and a mass range of 300-1500 m/z. A maximum injection time of 50 ms, normalized AGC target of 250% and RF Lens set to 30% was selected. Data was collected in centroid mode. MIPS (monoisotopic peak and peptide mode), Charge state (2-7), Precursor Selection Range (375-1500 m/z), Dynamic Exclusion(exclude after 1 time, 15 sec exclusion time, 10ppm low and high mass tolerance, excluding isotopes), Intensity Threshold (intensity range of 20000 (min) – 1E20 (max) with relative intensity threshold of 20%) filters were included. Data-dependent MS2 scans were acquired using cycle time (3 sec), a normalized HCD fragmentation energy of 30%, and a quadrupole isolation width of 0.7m/z. Fragments were detected using the Ion Trap with Turbo Mode at a scan range of 150-2000 m/z. A maximum injection time of 35 ms, normalized AGC target of 100%, and maximum injection time mode of dynamic was used. Data was also collected in centroid mode.

### 2.6 Database Search Parameters and Peptide Acceptance Criteria

Thermo .RAW files were processed using Spectromine (Version 4.4.240326) using 1% peptide and protein FDR search constraints, all settings were system defaults. Match Between Runs is not an option in Spectromine and therefore not implemented. The reference Homo sapiens protein .fasta from UNIPROT (201908, 98,300 entries). Variable modifications consisted of oxidation (M), Phosphorylation (S, T, Y), and protein N-terminal Acetylation. Carbamidomethylation was the sole static modification for the Trypsin search.

### 2.7 Data Analysis & Availability

Post-database searching analysis was performed as previously described^19,22^ using Python 3.12 with matplotlib (version 3.9.1), scipy (version 1.14.0), numpy (version 2.0.1), scikit-learn (version 1.5.1), pandas (version 2.2.2), py-venn, and a custom ‘helpers’ module.

For protein profiling samples, the Protein Groups, Peptides, and PTM reports were exported to .csv files and loaded into Python. For Protein Groups, Peptides, and PTMs, unique entries were determined by Gene Name, Stripped Sequence, or Flanking Sequence, respectively. For protein profiling PTMs, only PTMs annotated as ‘Phospho (STY)’ on the amino acid Y were kept. Potential duplicates were removed by keeping entries with the fewest missing values and highest median intensity, as previously described^22^. Sample counts were determined by counting the number of non-missing value entries in a label-free quantitation column for a particular sample. For venn diagrams, an entry was kept for a manufacturer/group if there was at least one non-missing value in that group for that entry. For pY samples, the Peptides and PTM reports were exported to .csv files and loaded into Python. Unique entries were determined equivalently as for protein profiling samples, except PTM samples were required to have the ‘Phospho (STY)’ modification only on Y, which is conservative in that only peptides where the most probable phosphorylatable amino acid is tyrosine are kept. Unique entry counting and venn diagrams were created as described for protein profiling samples. Statistical significance between stimulated and unstimulated samples for each pY site was determined using a Welch’s T-test and global false discovery rate adjustment using the method of Storey^25,26^. At no point during the analysis were missing values imputed. All code, graphs, and original, unprocessed Spectromine output files are available on GitHub (https://github.com/Aurdeegz/Desalting-Comparison/tree/master).

All of the post-searching data are available in the Supporting Information (Tables 1-5). Raw files are available from the ProteomeXchange Consortium via the Pride partner repository (Dataset ID: PXD064575).

### 2.8 Western blotting

Western blotting was performed as previously described^19,20^. Briefly, samples were separated on home-poured 10% polyacrylamide gels at 100 V for 120 minutes before transferring to a PVDF membrane (Millipore Sigma #IPFL00010) at 100 V for 100 minutes. Membranes were blocked using a TBS blocking buffer (LI-COR #927-60001) before overnight incubation in a primary antibody solution. The next morning, membranes were washed three times in 1X TBST before incubation in a secondary antibody solution for 60 minutes at room temperature. Finally, the membranes were washed three times in 1X TBST and twice in 1X TBS before imaging on a LI-COR Odyssey M scanner. Quantification was performed using LI-COR ImageStudio Version 5.2, and statistical analysis was performed in Python 3.12 as previously described^19^.

Primary antibodies: pY clone 4G10 (1:2000, Cell Signalling Technologies #96215), GAPDH (1:10000, Millipore Sigma #G9545), PLCγ1 (1:10000, Millipore Sigma #05-163), PLCγ1^Y783^ (1:2000, Cell Signalling Technologies #2821), Erk1/2 (Cell Signalling Technologies #9107), Erk^T202Y204^ (1:2000, Cell Signalling Technologies #9101)

Secondary antibodies: IRDye 680RD Goat anti-Mouse IgG (LI-COR #926-68070) IRDye 800CW Goat anti-Mouse IgG (LI-COR #926-33210), IRDye 680RD Donkey anti Rabbit IgG (LI-COR #926-68073), IRDye 800CW Donkey anti Rabbit IgG (LI-COR #926-32213)

## 3 Results

### 3.1 Experimental design and rationale

To test the capability of commercially available, high protein input desalting products for phosphoproteomics, we use a Jurkat T cell receptor (TCR) stimulation model, which has very well described tyrosine phosphorylation and RAS/MAPK activation patterns^27,28^ (Figure 1). Five replicates of basal state (0 minute) and stimulated (2 minutes with CD3/CD28 crosslinking) are first lysed in 8 M Urea, then split into 4.5 mg aliquots and volume normalized for input to desalting methods. For sample processing and/or desalting applications, we use ProtiFi S-Trap, a centrifugal sample preparation column, TECAN’s Narrow Bore Extraction (NBE), reverse-phase columns compatible with the Resolvex A200 liquid handling system for semi-automated desalting, Pierce C18 microcentrifuge columns, and Waters’ Sep-Pak C18 vacuum cartridges. After desalting, 1/100 of each sample (∼45 μg) was taken for protein profiling, and the remaining samples (∼4 mg) were used for phosphotyrosine (pY) enrichment using the Src SH2 superbinder (Figure 1A). Two minute antibody stimulation of TCR signalling is confirmed by α-pY and α-Erk1^T202Y204^/α-Erk2^T185Y187^ Western blot, which show a significant increase in pY and phosphorylated Erk abundance in the 2 minute stimulated samples compared to the 0 minute samples (Figure 1B, Supporting Figure 1).

### 3.2 Usability, peptide recovery, and cost for first-pass desalting products

Physical setups of each digestion platform are noted in Figure 2 A-D, with only Waters and TECAN requiring specialized equipment to operate them (a Vacuum Manifold and Resolvex A200 positive Pressure Manifold respectively). We first sought to qualitatively evaluate the usability, cost, and effectiveness of each first-pass desalting product. As shown in Figure 2E and F, TECAN NBE columns have the lowest hands on desalting time for any system (53 minutes desalting time, with ∼ 5 minutes hands on), whereas Pierce C18 columns are the most laborious overall (2 hours and 3 minutes for desalting) due to the screw-cap design. ProtiFi S-Trap columns have a much longer time before trypsin addition (3 hours 35 minutes) due to some slow flowing columns during the washing centrifugation steps, however, there is almost no time requirement following the addition of trypsin (20 minutes). Waters Sep-Pak C18 cartridges, the preferred desalting method in the Salomon laboratory, have a highly variable desalting time (46 minutes from an expert user, but ∼2 hours for new users) and a large elution volume (7 mL), requiring lyophilisation for drying. On average, S-Trap had the highest percent recovery, as measured by A205 NanoDrop (Figure 2G) of protein profiling samples, of any product (76.02%), followed by Pierce C18 columns (58.53%), Sep-Pak (40.52%), and NBE columns (28.06%). Higher recovery percentage from S-Trap comes with an increased cost, as S-Trap columns are also the most expensive at ∼$17.50 per sample for the smallest purchasable amount (Figure 2H). In summary, while TECAN NBE extraction is, by far, the easiest method requiring the least user interaction, ProtiFi S-Trap columns provide the highest percent recovery of any product tested.

**Figure 2.**
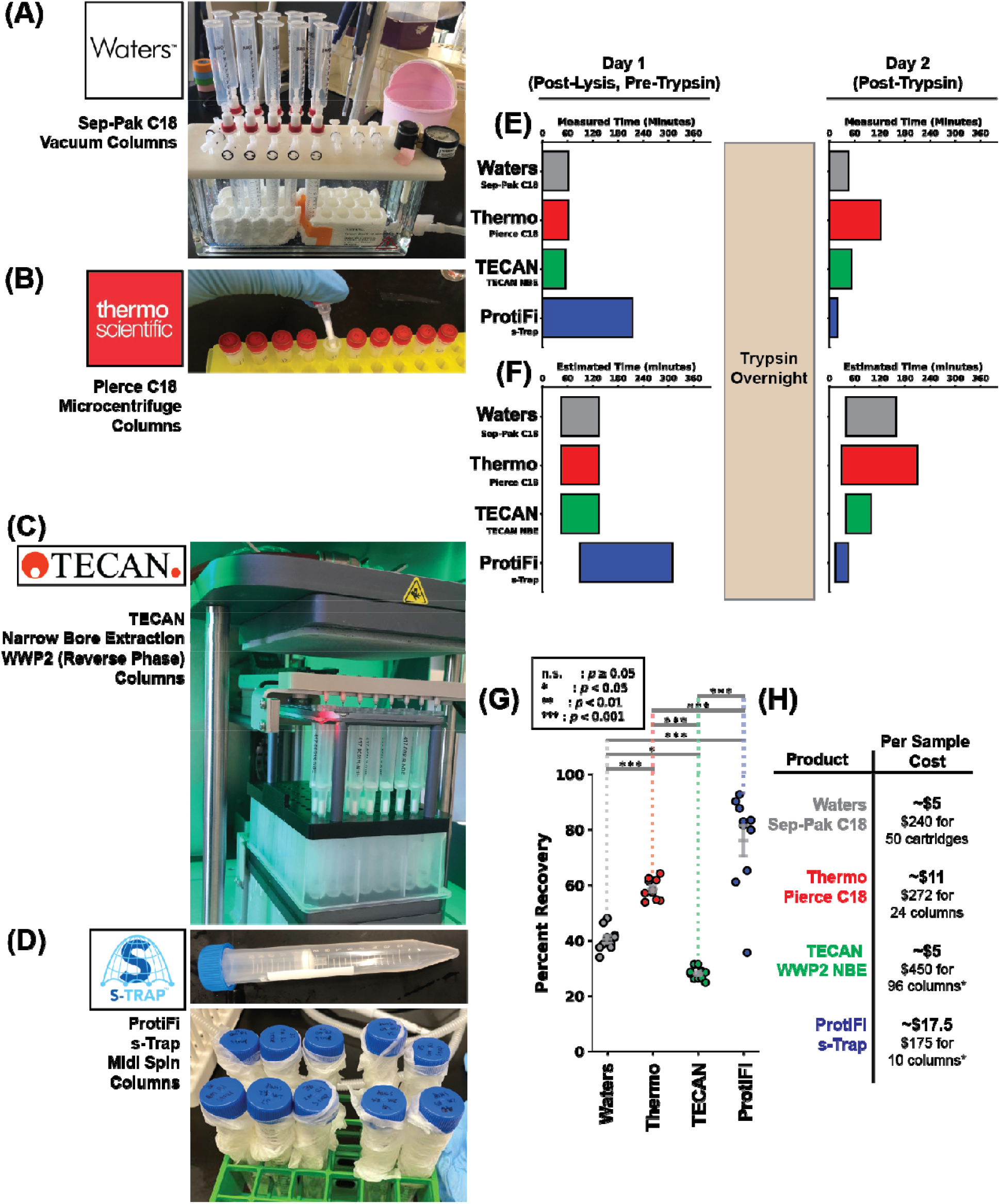
TECAN NBE columns provide superior reproducibility, automation capability, and cost efficiency, while ProtiFi S-Trap columns provide superior recovery. (A-D) Pictures showing Waters Sep-Pak C18 columns, Pierce C18 spin columns, TECAN NBE columns, and ProtiFi S-Trap columns, respectively. (E) Schematic showing measured time for post-lysis, pre-trypsin experimental time (left) and post-trypsin time (right). (F) As in (E), except showing Estimated Time. (G) Graph showing Percent Recovery of peptides after desalting using the four products. FWER adjusted p-values are determined by Fisher’s LSD corrected using the method of Holm and Sidak.

### 3.3 ProtiFi S-Trap provides superior peptide and protein identification

We find that, in line with the percent recovery, ProtiFi S-Trap columns provide the highest total ion current (TIC) with an average TIC max of 3.48E10, compared with 2.27E10 for Waters Sep-Pak, 2.31E10 for Pierce Spin Columns, and 2.32E10 for TECAN NBE columns (Figure 3A, Supporting Figure 2). Notably, TECAN NBE columns, across all replicates, significantly undersampled peptides that typically elute in the aqueous portion of the gradient(<20%Solvent B) as the TIC indicates slightly more hydrophobic peptides eluting at approximately 30-40 minutes into the gradient (i.e >25% Solvent B) compared to ∼10-20 minutes for all other products (Figure 3A, Supporting Figure 2). Principal component analysis of stripped peptide sequences shows replicate clustering for each product with Waters Sep-Pak and Pierce columns clustering together. Multiple regression shows that TECAN NBE columns have the highest reproducibility, with a correlation coefficient of R=0.919 (Figure 3B). ProtiFi S-Trap has significantly higher peptide sequencing of all the products, whereas TECAN NBE columns have the lowest stripped sequence count (Figure 3C). Of the 54,869 unique stripped peptide sequences we identify in S-Trap samples, 16,740 sequences are observed using all products and 25,654 are uniquely sequenced using S-Trap (Figure 3D). Evaluating protein groups, we see similar trends. S-Trap and TECAN cluster separately and Waters Sep-Pak and Thermo columns cluster together (Figure 3F). For protein groups, Pierce C18 spin columns have the highest replicate reproducibility with a correlation coefficient of R=0.942 (Figure 3D). ProtiFi S-Trap identifies the most unique protein groups and TECAN NBE columns identify the least (Figure 3E). While 3,808 protein groups are identified amongst all the products, ProtiFi S-Trap identifies 375 unique protein groups compared with 1 for TECAN NBE columns, 0 for Pierce columns, and 1 for Waters Sep-Pak columns (Figure 3G). In agreement with the elevated sequencing of the proteome, S-Trap samples also show markedly improved sequencing of the phosphoproteome (pS, T, Y) in whole cell digest samples, sequencing about 2-fold more phosphorylation sites per replicate with moderate reproducibility and identifying 483 unique phosphorylation sites compared with the other products (Supporting Figure 3). Together, our data suggest that, when using a standardized workflow for protein profiling, ProtiFi S-Trap columns provide superior sequencing into the proteome of human T cells.

**Figure 3.**
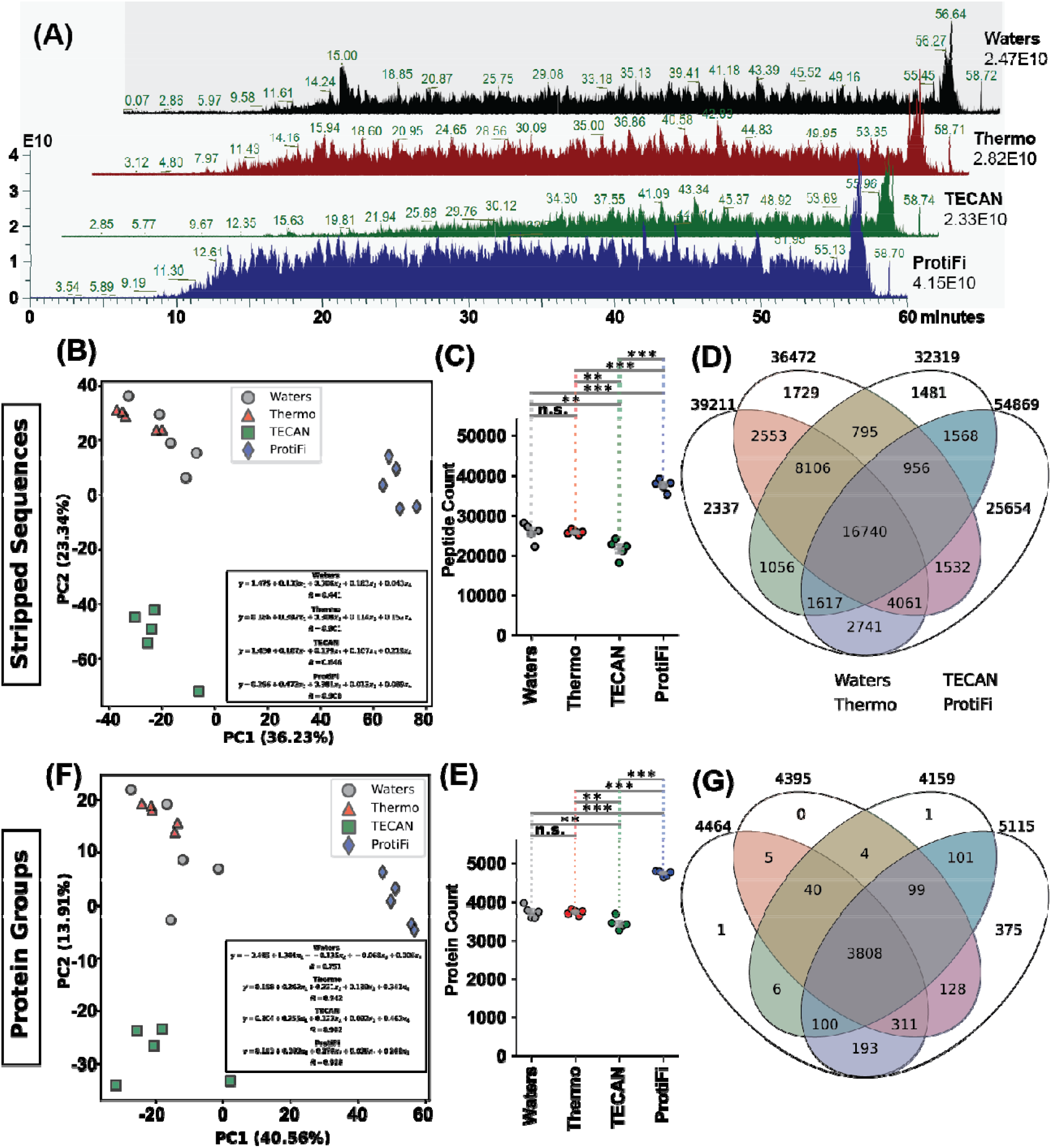
ProtiFi S-Trap provides superior peptide and protein identification by LC-MS. (A) TIC for Replicate 1 whole cell digest samples. Labels to the right show the global maximum ion current for the indicated sample. (B) Principal component analysis of whole cell digest samples at the stripped peptide sequence level. Inset shows the line of best fit but multiple regression. (C) Graph showing sample-specific counts of stripped peptide sequences. P-values are determined as indicated in Figure 2G. (D) Venn-diagram showing the overlap in stripped peptide sequence identification between the tested products. (F-G) As in B-D), except protein groups are used as input.

### 3.4 ProtiFi S-Trap provides superior pY site identification

We next sought to determine whether specific first-pass desalting products improved the efficacy of a well-characterised pY enrichment protocol whereby phosphotyrosine samples are acquired by performing Src SH2 superbinder pY enrichments. After generating lyophilized enriched samples, peptides are reconstituted in 35 μL of 5% Acetonitrile and 0.1% Formic Acid, and injecting 15 μL (∼43% of the total sample) due to the low abundance of tyrosine phosphorylation in cells^29–31^. In contrast to protein profiling samples, the TIC shape for pY samples appears to correlate with the choice of manufacturer but does not qualitatively correlate with percent recovery. Samples prepared using Waters Sep-Pak C18 columns appear to have highly abundant, hydrophilic peptide species that do not appear with any other vendor. Further, ProtiFi S-Trap samples appear to have the lowest, most stable TIC intensity throughout the run, whereas Pierce C18 spin columns and TECAN NBE columns have a similarly organic-heavy (i.e hydrophobic peptide rich) TIC pattern (Figure 4, Supporting Figure 4). Unique pY sites sequenced in all 0 minute samples are highly reproducible irrespective of the product used, with replicates generally clustering by PCA and correlation coefficients > 0.9 (Figure 4B, Supporting Figure 5). Although variable, ProtiFi S-Trap outperformed all other products in pY site sequencing with between 300 and 550 pY sites per replicate (Figure 4C) and 415 pY sites uniquely identified in at least one S-Trap sample (Figure 4D). These trends hold for α-CD3ε/α-CD28 stimulated samples as well; samples prepared using S-Trap are highly reproducible and sequence the tyrosine phosphoproteome much deeper than any other tested product (Figures 4F-G). Samples prepared using S-Trap have the highest unphosphorylated peptide sequencing among the tested products and Waters Sep-Pak samples have the lowest, although unphosphorylated peptide sequencing appears to be much less reproducible with correlation coefficients generally 0.7 or less (Supporting Figures 6 & 7). Together, these results show that samples prepared using ProtiFi S-Trap before pY enrichment sequence the pY proteome deeper than samples prepared with Waters Sep-Pak C18 cartridges, Pierce C18 spin columns, or TECAN NBE columns.

**Figure 4.**
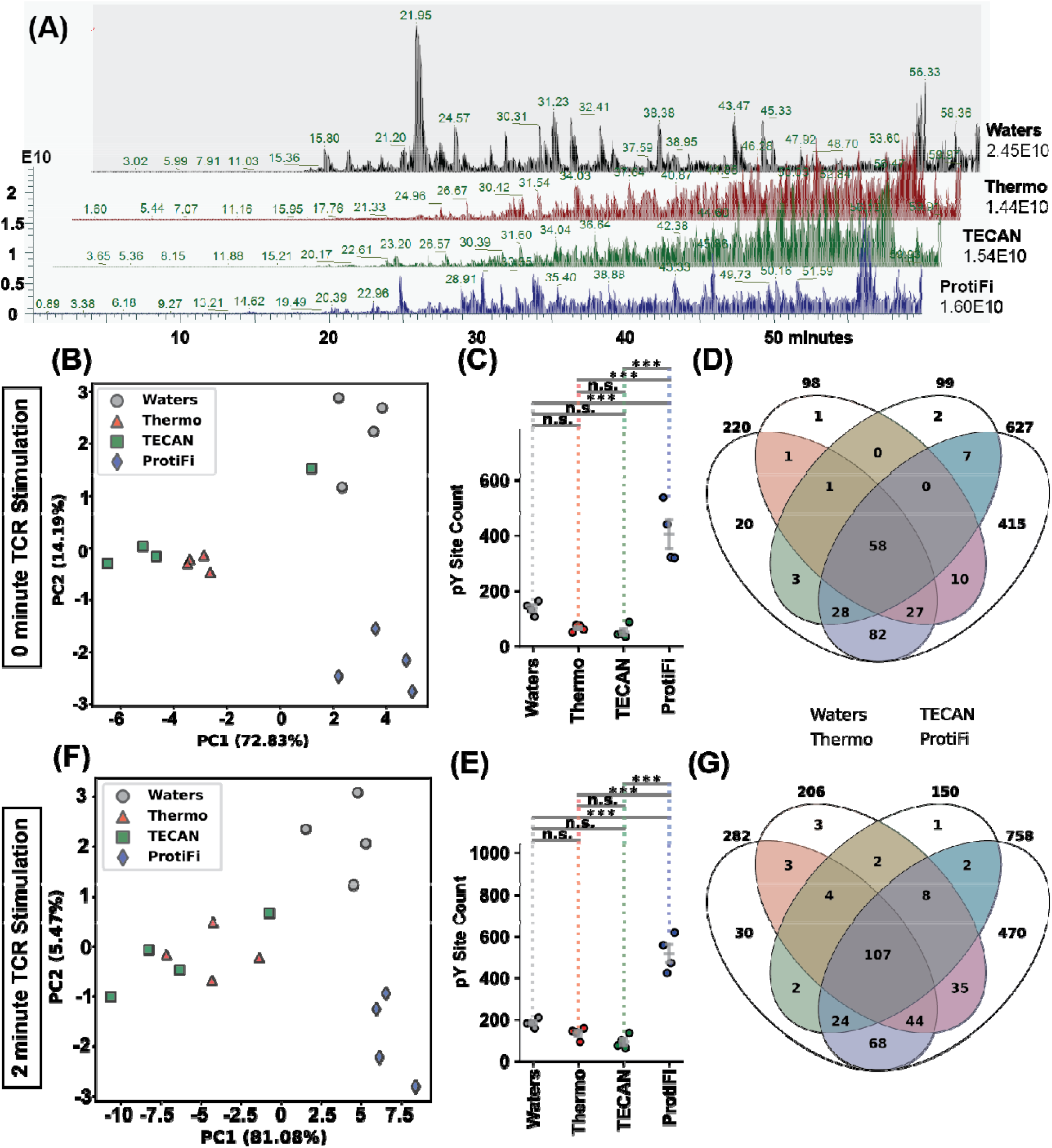
Samples prepared with ProtiFi S-Trap show reproducible and superior pY sequencing. (A) Total ion current for 0 minute Replicate 1 pY enrichment samples. (B) Principal component analysis of 0 minute samples at the unique pY site level. (C) Graph showing sample-specific counts of unique pY sites. P-values are determined as indicated in Figure 2G. (D) Venn-diagram showing the overlap in unique pY site identification between the tested products. (F-G) As in B-D), except 2 minute stimulation samples are used as input.

**Figure 5.**
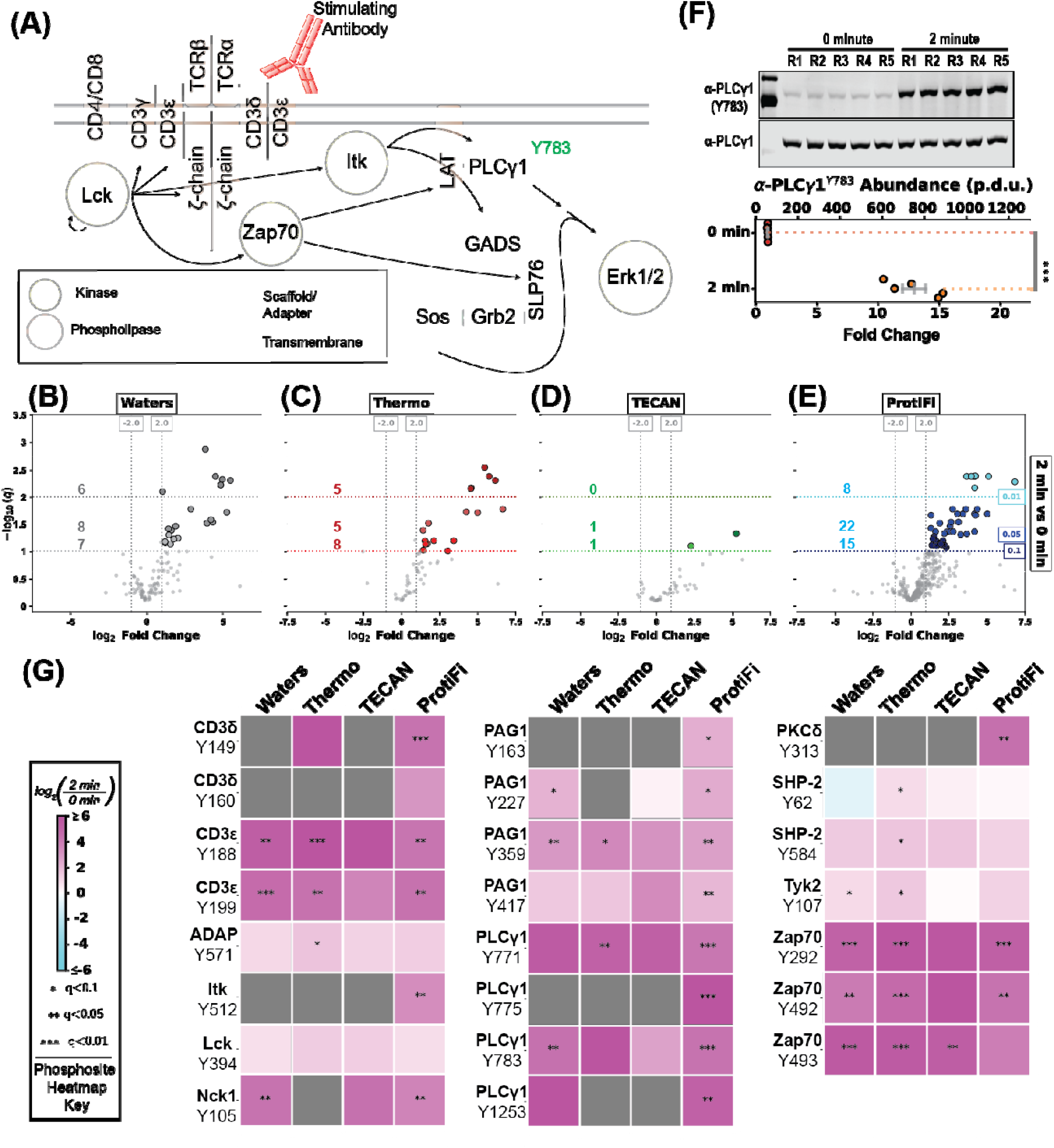
Samples prepared using ProtiFi S-Trap show superior identification of biologically relevant, T cell receptor responsive pY sites. (A) Simplified schematic of T cell receptor signalling. (B-E) Volcano plot analysis comparing pY site abundance between 2 minutes and 0 minutes of stimulation for samples prepared using Waters Sep-Pak C18 columns, Pierce C18 spin columns, TECAN WWP2 NBE columns, and ProtiFi S-Trap spin columns, respectively. Values to the left indicate the number of unique pY sites in a given threshold (0.1 > q ≥ 0.05, 0.05 > q ≥ 0.01, 0.01 > q from bottom to top). (F) Western blot analysis of PLCγ1^Y783^ with accompanying quantification. ^***^ indicates p < 0.001 by a Welch’s T-test. (G) Heatmaps comparing 2 minutes and 0 minutes of T cell receptor stimulation for unique pY sites. Grey squares indicate n ≤ 3 replicates for a given pY site in either 2- or 0-minute groups. ^*^ indicates 0.1 > q ≥ 0.05, ^**^ indicates 0.05 > q ≥ 0.01, ^***^ indicates 0.01 > q.

### 3.5 ProtiFi S-Trap provides superior pY site changes amongst desalting products

To evaluate whether the increase in sequencing depth corresponds to an increase in biologically meaningful information, we compare the 0 minute and 2 minute α-CD3ε/α-CD28 stimulated samples and evaluate the phosphorylation of key T cell receptor signalling proteins (Figure 5A). When comparing all sample preparation methods, we observed an increase in pY abundance with Waters Sep-Pak, Pierce C18, and ProtiFi S-Trap showing 14, 10, and 30, significantly changing unique pY sites respectively, (q<0.05; Figure 5B, 5C, & 5E), in line with our Western blot data showing high pY induction (Figure 1B). Samples prepared using TECAN WWP2 NBE columns show a single significantly changing pY site, although other sites appear to be increased albeit not significantly (Figure 5D). By Western blot analysis, we observed a significant increase in PLCγ1^Y783^, the activation site on PLCγ1 and a substrate of interleukin-2 inducible T cell kinase (Itk)^32^, which is also significantly increased in ProtiFi S-Trap and Waters Sep-Pak sample preparations (Figures 5F-G). As expected, samples prepared using ProtiFi S-Trap columns show significant induction of many known T cell signalling phosphorylation sites, including the immunotyrosine based activation motif (ITAM) sites CD3ε^Y188^ and CD3ε^Y199^, Nck^Y105^, PAG1^Y417^, PLCγ1^Y771^, Zap70^Y292^, and Zap70^Y492^, all of which are also seen significantly changing when using at least one other product. Some important phosphorylation sites, including the ITAM CD3δ^Y149^, the activation site Itk^Y512^, the inhibitor site PLCγ1^Y775^, and the regulatory sites PLCγ1^Y1253^ and PKCδ^Y313^ are exclusively seen as significantly changing when using ProtiFi S-Trap for sample preparation. Many other phosphorylation sites are also seen trending towards significance (0.05 ≤ q < 0.1) including ADAP^Y571^, SHP-2^Y62^, SHP-2^Y584^ and Tyk2^Y107^ in Pierce C18 preparations (8 total), PAG1^Y227^ and Tyk2^Y107^ in Waters preparations (7 total), and PAG1^Y163^ and PAG1^Y227^ in ProtiFi preparations (15 total; Figures 5B, 5C, 5E, & 5G). In summary, our data show that increased pY sequencing corresponds to an increase in unique, significantly changing, and biologically meaningful pY sites observed by LC-MS.

## 4 Discussion

The sensitivity of modern mass spectrometers allows for the quantification and unique identification of thousands of peptides, with applications spanning from identifying proteins within single cells, non-denatured or intact proteins for structure-based analysis, and post-translational modification mapping^2–6,8,10,11^. Further, automation using liquid handling systems has profoundly improved reproducibility of proteomics sample analysis and lowered the barrier to entry for new proteomics users^16–18^. Salt removal is critical for the preparation of all proteomics samples, as residual salts and detergents interfere with peptide ionization and ultimately detection by MS^15^. Research laboratories and core facilities have an array of commercial desalting products to choose from, and frequently choose a product based on their training, specific applications, or automation readiness. While evaluations of low protein input desalting products exist^15,33^, currently no work evaluating high protein input desalting products for applications to pY proteomics exists to our knowledge.

Here, we provide the first evaluation of high protein input desalting methods for use in pY proteomics applications. Using four commercially available desalting products - Waters Sep-Pak C18 vacuum cartridges, Pierce C18 spin columns, TECAN WWP2 narrow bore extraction columns, and ProtiFi S-Trap spin columns - we evaluate the usability, recovery, peptide/protein profiling, and pY sequencing using the Jurkat T cell receptor signalling model (Figure 1), which has very well-established tyrosine phosphorylation patterns^27^. Waters Sep-Pak columns on a low vacuum manifold require patience, focus, and good organizational skills, but are overall not very time consuming (Figure 2A, E-F). Pierce spin columns are the most laborious to use due to their low volume format (∼300 μL) and the relatively high volumes (3-10 mL) required for dissolving and digesting 4 mg of protein (Figure 2B, E-F). Using TECAN NBE columns, which may also be used for on column digestion (not evaluated here) on the Resolvex A200 liquid handling system has nearly no hands on time for desalting, is procedurally very simple once a method is developed, and is very reproducible (Figure 2C, E-F). ProtiFi S-Trap which requires manual operation and a centrifuge, is unique in comparison to other products evaluated here in that the platform performs both desalting and digestion in suspension on the column. While this increases the hands-on time for pre-digestion processing, it actually reduces the time post-digestion substantially (Figure 2D-F). Of all the desalting methods, ProtiFi S-Trap had the highest recovery variability and yet highest overall percent recovery (∼75% on average), while TECAN NBE columns had the lowest recovery (∼30%) yet highest reproducibility. Thus, a Resolvex A200 compatible format that combines the benefits of automation and reproducibility desired for high-throughput proteomics workflows, with ProtiFi S-Trap spin column chemistry may provide the most complete peptide recovery for high protein input digestion.

We find that higher peptide recovery translates to increased peptide, protein group, and pY site identification using a standardized LC-MS workflow. While Waters Sep-Pak and Pierce spin columns had similar recovery (∼40-60%), reproducibility (R>0.88), and peptide/protein identifications (∼26,000 peptide, ∼3,800 protein), TECAN WWP2 NBE columns had fewer identifications (∼22,000 peptide, 3,300 protein), likely due to low recovery of hydrophilic peptides (Figure 3). ProtiFi S-Trap, in contrast, vastly outperformed all other desalting methods, identifying ∼37,000 unique peptide sequences corresponding to ∼4,600 protein groups, with 25,748 uniquely identified peptides and 384 uniquely identified proteins (Figure 3). The increase in sequencing depth using ProtiFi S-Trap for first-pass sample preparation corresponds to an increase in repeatable pY site identification, sequencing about 300-550 pY sites per replicate and identifying more than 400 unique pY sites (Figure 4). Further, the increase in unique pY sites observed using ProtiFi S-Trap corresponds to an increase in TCR responsive pY sites, including many biologically meaningful pY sites on key TCR signalling proteins (Figure 5).

Surprisingly, we did not solubilize S-Trap input proteins in SDS, as required by the manufacturer^34^, yet the S-Trap columns still outperformed traditional and novel desalting methods.

With an increasing interest in proteomics applications and the astounding variability in sample preparation techniques, having automatable, reproducible workflows is necessary to meet demand and expectations. The TECAN Resolvex A200 provides a reliable, easy-to-use liquid handling system suitable for high protein input desalting applications, however ProtiFi S-Trap chemistry provides superior recovery and identifications by LC-MS. For low-protein input (<300 ug) applications, ProtiFi sells a Resolvex A200 liquid handling system compatible version of S-Trap, and our results indicate that a high-protein input product would be ideal for PTM proteomics applications that require high protein input amounts.

## 5 Associated Content

Supporting information are available free of charge, including

- (.DOCX) All figures in the supporting information, including Western blots of T cell activation, TIC for protein profiling and pY enriched samples, regression analysis of reproducibility, stripped sequences identified in pY enriched samples.
- (ZIP) Excel tables showing formatted output of statistical analysis for protein profiling samples (Tables 1-3) and pY samples (Tables 4 & 5).

## Acknowledgements

This research is based in part upon work conducted using the Proteomics Core Facility, which was supported in part by the National Institutes of Health Grant No. 1S10RR027027 (Orbitrap XL-ETD), 1S10OD036295 (Ascend Tribrid, FAIMS, Vanquish Neo), and the Division of Biology and Medicine, Brown University. We would also like to acknowledge NIH grant P01AI091580.

