## Supplementary material for "Evaluating First-Pass, High Protein Capacity Desalting Techniques For Phosphoproteomics Applications": supporting_information.docx

**Graphical Abstract:**

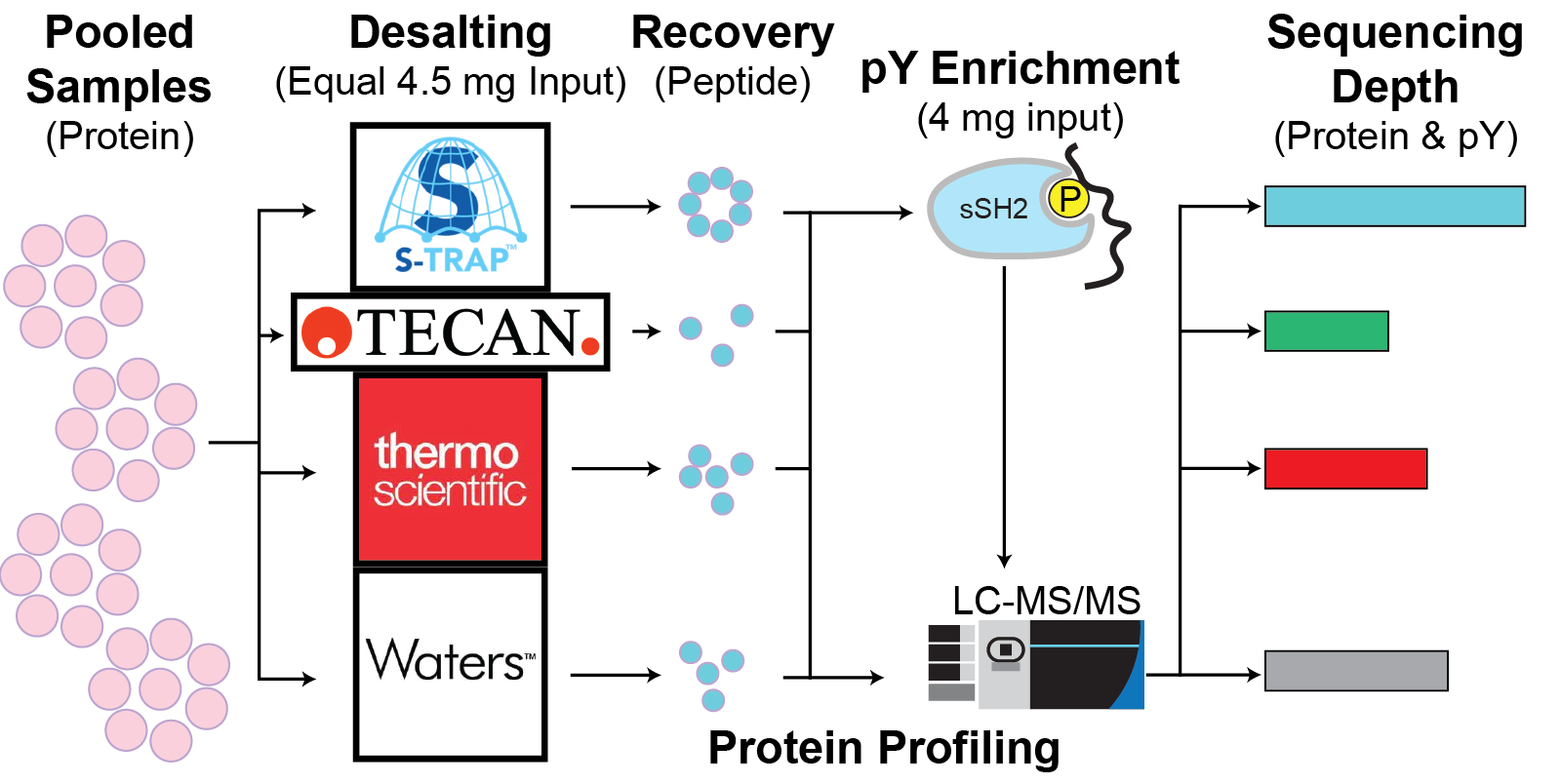

**Abstract:**

Many commercial desalting products exist for pre-MS peptide cleanup, although few exist that can handle the high protein input (≥ 4 mg) required for phosphotyrosine enrichment. For these desalting products, the technical aptitude required for effective and organized desalting is often a barrier to entry for new users. Here, we evaluate four commercially available desalting techniques with varying degrees of automation, operational organization, and chemistries to determine the most cost-effective, user-friendly, and sensitive technique for protein profiling and phosphotyrosine (pY) enrichment. We find that TECAN Narrow Bore Extraction (NBE) products are the most cost effective per sample and least difficult to use, whereas ProtiFi’s S-Trap are the most expensive per sample and Pierce C18 spin columns have the worst operational organization. ProtiFi S-Trap vastly outperforms other desalting methods for peptide sequencing and protein profiling applications, uniquely identifying 25,654 unique peptide sequences and 375 unique proteins. Consistently, ProtiFi S-Trap samples show the deepest pY sequencing after Src SH2 superbinder enrichment, leading to the highest identification of significantly changing, biologically relevant pY sites in a Jurkat T cell signalling model. Our data show that ProtiFi S-Trap columns provide high peptide recovery, thus increasing meaningful pY site identification.

### **Table of Contents**

**Supporting Table 1**: (.XLSX) Complete list of unique proteins identified in WCD samples

**Supporting Table 2**: (.XLSX) Complete list of unique Phospho (STY) sites identified in WCD
samples

**Supporting Table 3**: (.XLSX) Complete list of unique stripped sequences identified in WCD samples

**Supporting Table 4**: (.XLSX) Complete list of unique Phospho (Y) sites identified in sSH2 enrichment samples

**Supporting Table 5**: (.XLSX) Complete list of unique stripped sequences identified in sSH2 enrichment samples

**Supporting Figure 1**: Western blot analysis of Erk1 and Erk2 phosphorylation

**Supporting Figure 2**: Total ion current for WCD samples

**Supporting Figure 3**: Analysis of Phospho (STY) sites observed in unenriched WCD samples

**Supporting Figure 4**: Total ion current for all 2 minute sSH2 enrichment samples

**Supporting Figure 5**: Multiple linear regression on unique pY sites from sSH2 enrichment samples

**Supporting Figure 6**: Analysis of unphosphorylated stripped sequences identified in 0 minute sSH2 enrichment samples

**Supporting Figure 7**: Analysis of unphosphorylated stripped sequences identified in 2 minute sSH2 enrichment samples

**Supporting Table 1:** Complete list of unique proteins identified in all whole cell digest samples. Each row contains a unique protein (ID column = Gene Name), as well as the label free quantitation, log transformation, and basic statistics (mean, standard deviation, CV%) for each unique protein.

**Supporting Table 2:** Complete list of unique Phosphorylation sites (STY) identified in all whole cell digest samples. Each row contains a unique phosphorylation site (ID column = Flanking sequence), as well as the label free quantitation, log transformation, and basic statistics (mean, standard deviation, CV%) for each unique phosphorylation site.

**Supporting Table 3:** Complete list of unique stripped peptide sequences identified in all whole cell digest samples. Each row contains a unique peptide (ID column = Sequence), as well as the label free quantitation, log transformation, and basic statistics (mean, standard deviation, CV%) for each unique peptide.

**Supporting Table 4:** Complete list of unique Phosphorylation sites (Y) identified in all sSH2 enriched samples. Each row contains a unique pY site(ID column = Flanking sequence), as well as the label free quantitation, log transformation, and basic statistics (mean, standard deviation, CV%) for each unique protein. Included also are the results of hypothesis testing (uncorrected Welch’s T-test and Storey q-value correction) between 2 minute and 0 minute stimulation samples per vendor.

**Supporting Table 5:** Complete list of unique stripped peptide sequences identified in all sSH2 enriched samples. Each row contains a unique peptide (ID column = Sequence), as well as the label free quantitation, log transformation, and basic statistics (mean, standard deviation, CV%) for each unique peptide.

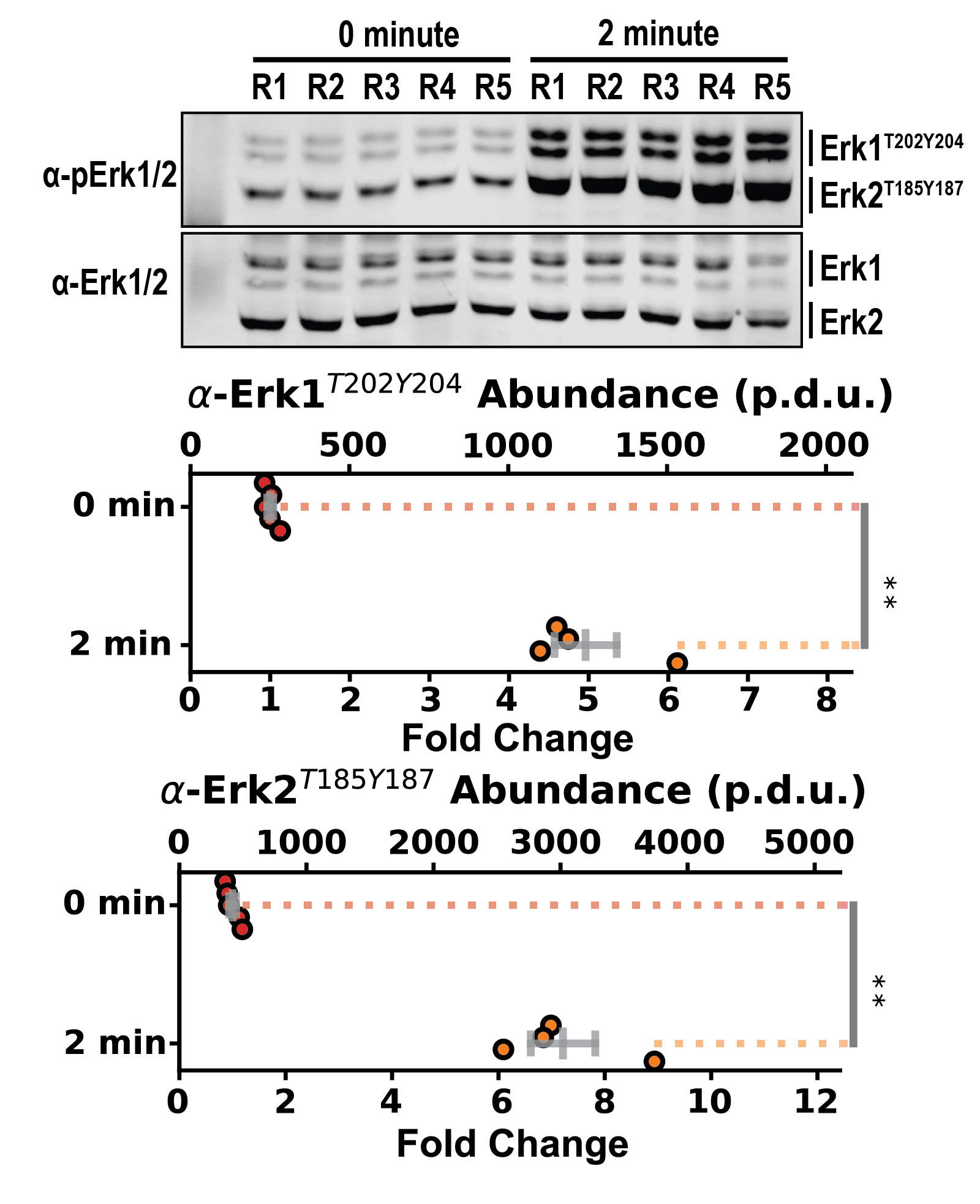

**Supporting Figure 1:** Stimulating antibodies lead to phosphorylation of Erk1^T202Y204^ and Erk2^T185Y187^. Western blot analysis of Erk1/2 phosphorylation, with accompanying quantification. ** indicates p < 0.01 by Welch’s T-test.

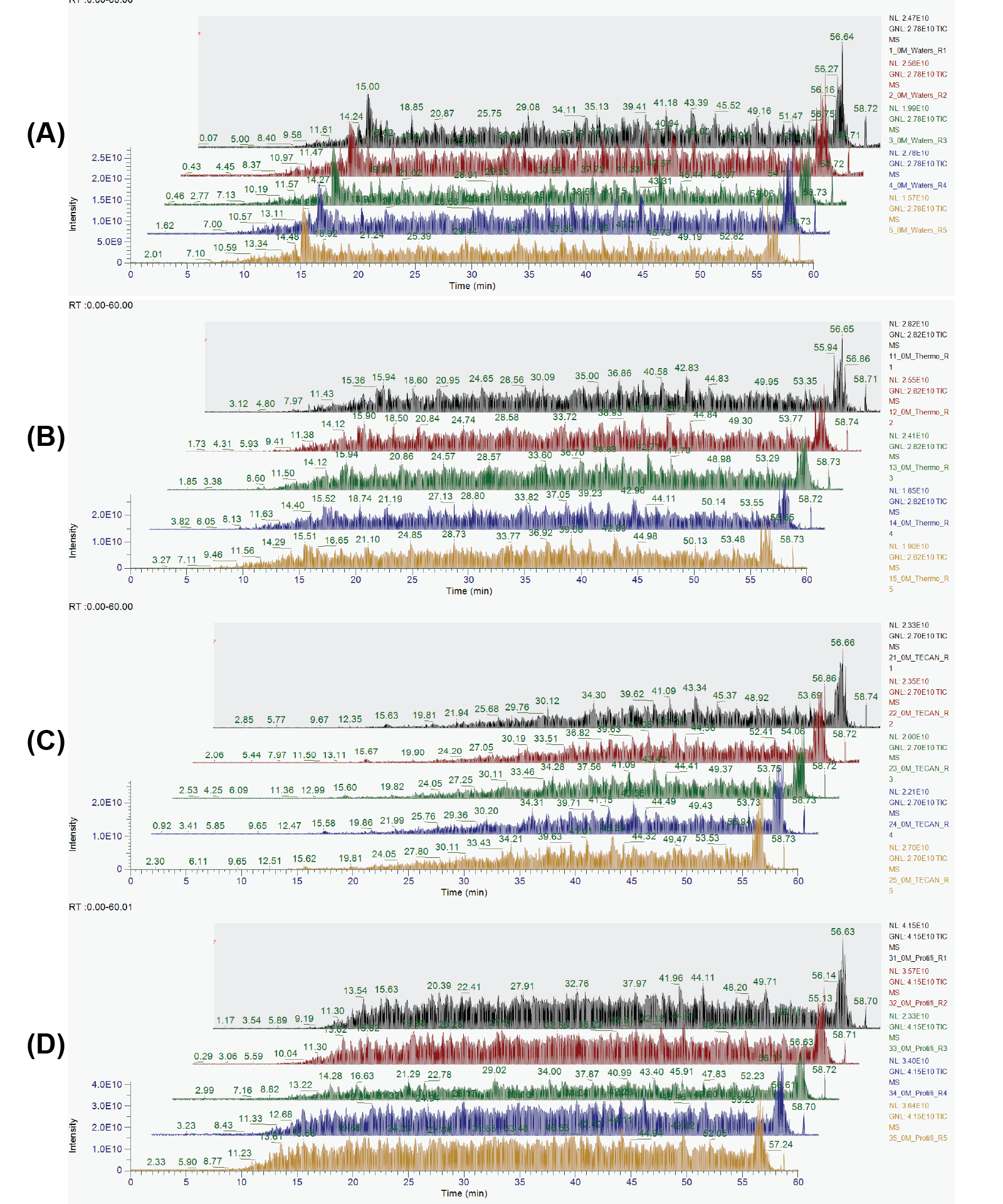

**Supporting Figure 2:** Total ion current for all whole cell digest samples from (A) Waters Sep-Pak vacuum columns, (B) Pierce C18 spin columns, (C) TECAN WWP2 Narrow Bore Extraction columns, or (D) ProtiFi S-Trap spin columns.

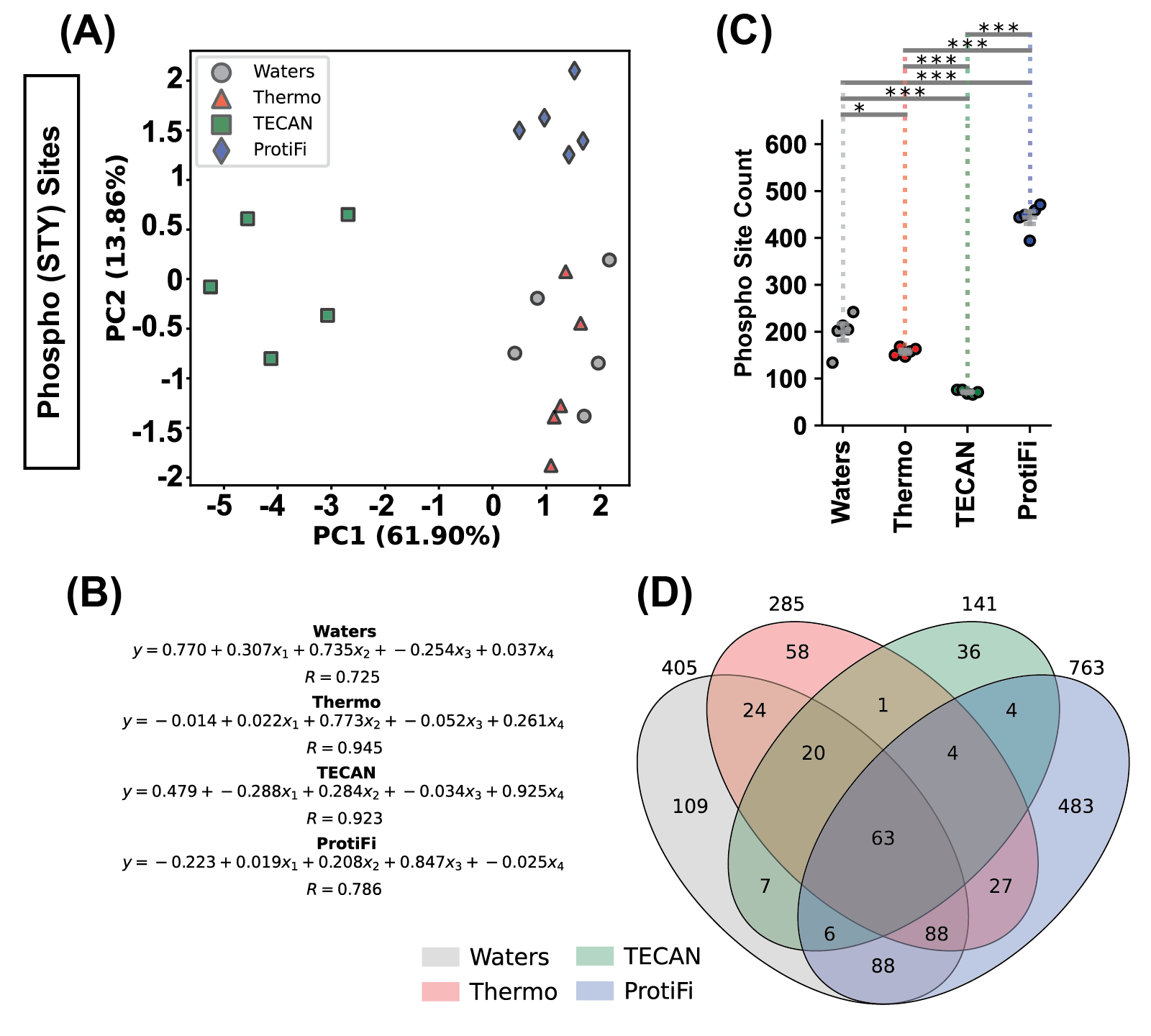

**Supporting Figure 3:** ProtiFi’s S-Trap sequences the phosphoproteome in protein profiling samples to a greater degree than other products. (A) Principal component analysis of Phospho (STY) Sites-containing peptides. (B) Multiple linear regression analysis on phosphorylation sites measured in all samples per manufacturer. (C) Graph showing sample-specific counts of unique phosphorylation sites. FWER adjusted p-values are determined by Fisher’s LSD corrected using the method of Holm and Sidak. * indicates p < 0.05, ** indicates p < 0.01, *** indicates p < 0.001. (D) Venn-diagram showing the overlap in unique phosphorylation site identification between the tested products.

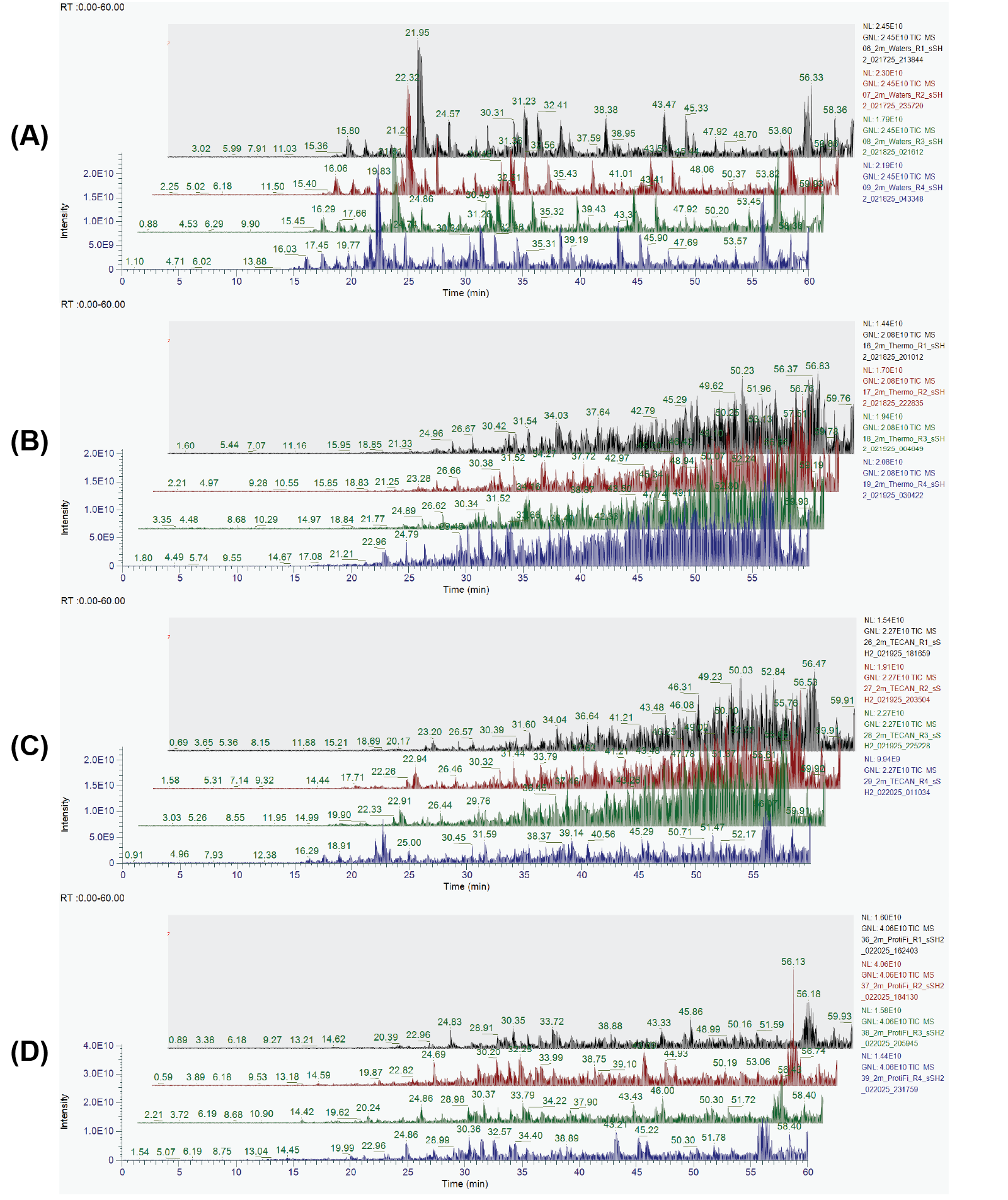

**Supporting Figure 4:** Total ion current for all 2 minute α-CD3ε/α-CD28 stimulation samples processed using (A) Waters Sep-Pak vacuum columns, (B) Pierce C18 spin columns, (C) TECAN WWP2 Narrow Bore Extraction columns, or (D) ProtiFi S-Trap spin columns.

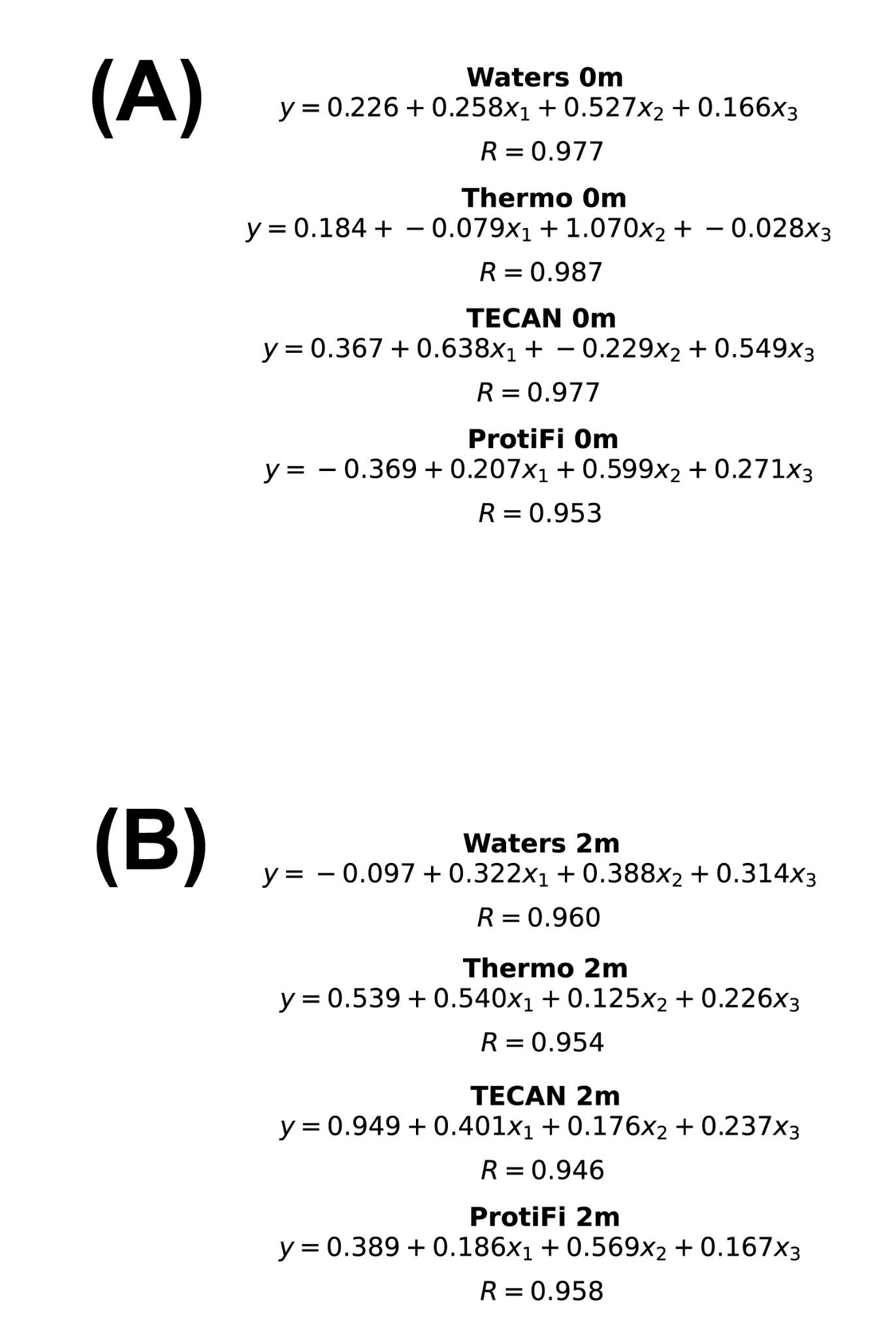

**Supporting Figure 5:** Replicate reproducibility as measured by Multiple Linear Regression for (A) 0 minute and (B) 2 minute α-CD3ε/α-CD28 stimulation samples

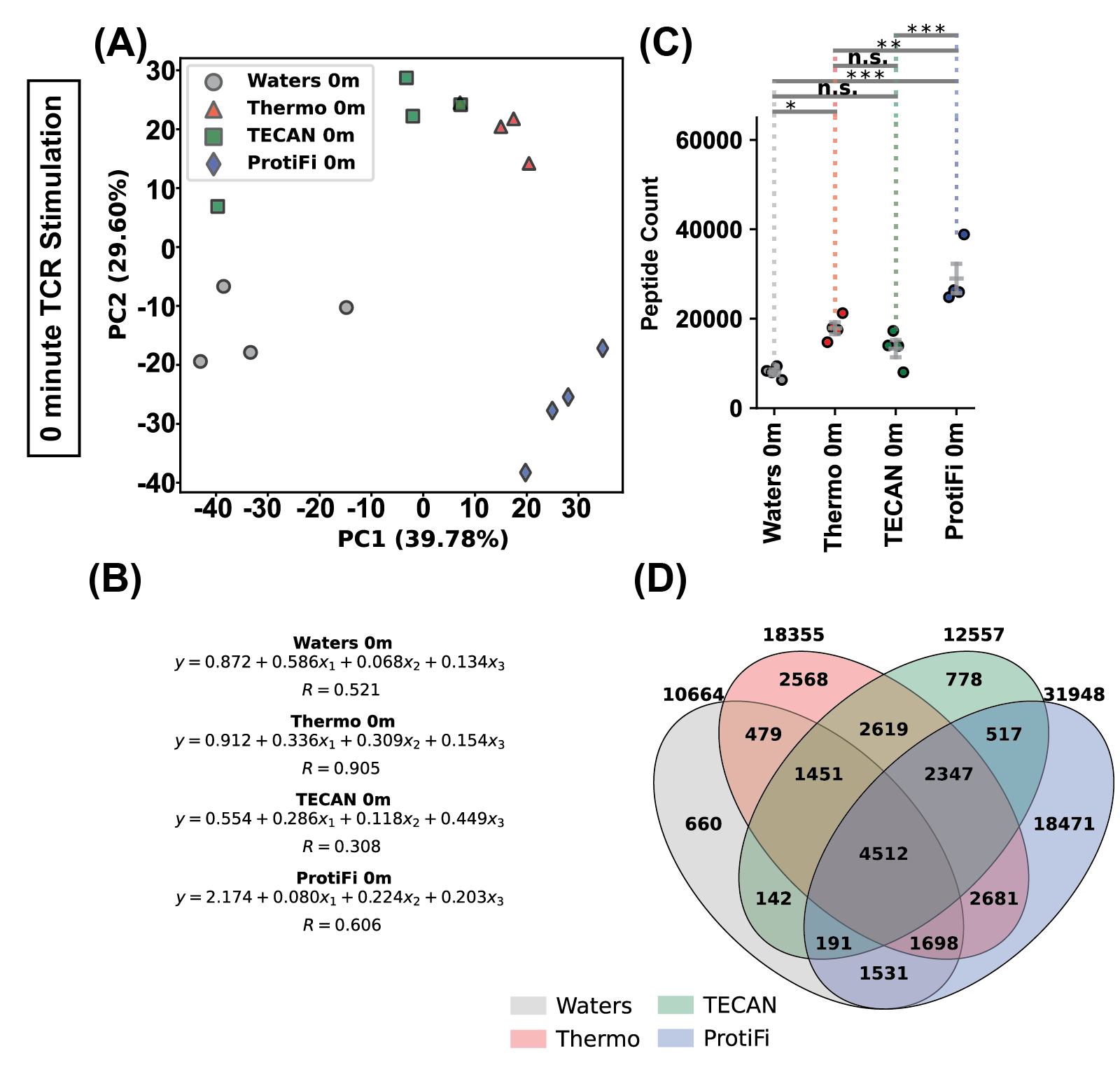

**Supporting Figure 6:** Background binding to sSH2 beads in unstimulated samples is highest when using ProtiFi’s S-Trap. (A) Principal component analysis of stripped sequences from sSH2-enriched samples. (B) Multiple linear regression analysis on stripped sequences measured in all samples per manufacturer. (C) Graph showing sample-specific counts of stripped sequences. FWER adjusted p-values are determined by Fisher’s LSD corrected using the method of Holm and Sidak. * indicates p < 0.05, ** indicates p < 0.01, *** indicates p < 0.001. (D) Venn-diagram showing the overlap in stripped sequence identification between the tested products.

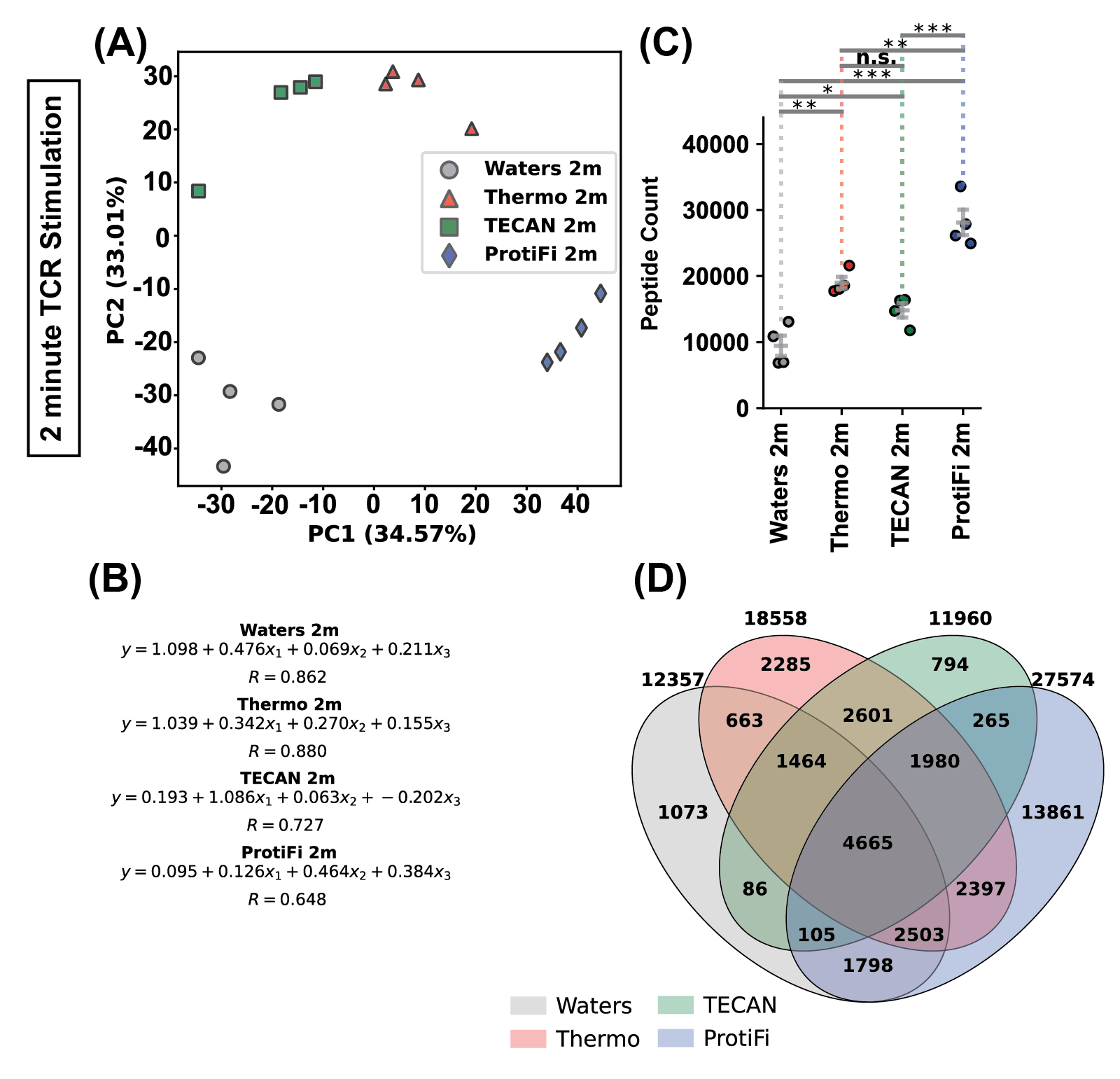

**Supporting Figure 7:** Background binding to sSH2 beads in stimulated samples is highest when using ProtiFi’s S-Trap. (A) Principal component analysis of stripped sequences from sSH2-enriched samples. (B) Multiple linear regression analysis on stripped sequences measured in all samples per manufacturer. (C) Graph showing sample-specific counts of stripped sequences. FWER adjusted p-values are determined by Fisher’s LSD corrected using the method of Holm and Sidak. * indicates p < 0.05, ** indicates p < 0.01, *** indicates p < 0.001. (D) Venn-diagram showing the overlap in stripped sequence identification between the tested products.
